# Kinetics of charged polymer collapse: effects of additional salt

**DOI:** 10.1101/2020.03.06.981431

**Authors:** Susmita Ghosh, Satyavani Vemparala

**Affiliations:** The Institute of Mathematical Sciences, C.I.T. Campus, Taramani, Chennai 600113, India; Homi Bhabha National Institute, Training School Complex, Anushakti Nagar, Mumbai, 400094, India

## Abstract

Extensive molecular dynamics simulations, using simple charged polymer models, have been employed to probe the kinetics and dynamics of early-stage collapse of charged polymers and the effect of additional monovalent salt on such kinetics. The exponents characterizing the coarsening dynamics during such early-collapse stage via finite size scaling for the case of charged polymers are found to be different from the neutral polymers, suggesting that the collapse kinetics of charged polymers are inherently different from that neutral polymers. The kinetics of coarsening of the clusters along the collapsed trajectory also depends significantly on the counterion valency and for higher valency counterions, multiple regimes are observed and unlike the neutral polymer case, the collapse kinetics are a function of charge density along the charged polymer. Inclusion of additional salt affects the kinetics and conformational landscape along the collapse trajectory. Addition of salt increases the value of critical charge density required to initiate collapse for all the counterion valencies, though the effect is more pronounced for monovalent counterion systems. The addition of salt significantly affects the collapse trajectory in the presence of trivalent counterions via promotion of transient long-distance loop structures inducing a parallel and hierarchical local collapsed conformation leading to faster global collapsed states. This may play a role in understanding the fast folding rates of biopolymers such as proteins and RNA from extended state to a collapsed state in the presence of multivalent counterions before reorganizing into a native fold.

## I. INTRODUCTION

Understanding the dynamics and kinetics along the collapsed trajectory of charged polymers underpins a variety of phenomenon such as protein folding, compact arrangement of DNA inside cells^1–5^ etc., It has been suggested that the biopolymers such as proteins and RNA fold via a fast collapse regime, followed by local rearrangement towards native state^6^. Single stranded molecules like RNA, which are strongly charged molecules, have been shown to form collapsed, albiet disordered, intermediate structures^7,8^. Such counterintuitive attractive electrostatic interactions between the like-charge moieties has been explained by three different models: counterion condensation ^9,10^, induced dipole effects^11^, and hydration effects^12^. The folding of RNA, in the presence of multivalent counterions, has been suggested to involve long-range tertiary interactions between parts of RNA which are distant from each other leading to transitory loop formation followed by closure of such loops and occurs on a milli-second time scale^6–8,13,14^. The compact (or collapsed) intermediate states provide a context in which such tertiary interactions can emerge and polyelectrolytes (PE)s have been long used as model systems to understand these diverse issues of collapse of such charged polymers. The collapse of charged polymers is also interesting to probe from a theoretical point of view, due to development of effective attractive interactions in the system, which is inherently a similarly charged system. Collapsed conformations of a neutral polymer in poor solvents is fairly well understood via the dominance of favorable excluded volume interactions between monomers of polymers^15–17^ leading to collapsed and extended conformations under poor and good solvent conditions. However, charged polymers have been shown to have collapsed conformations in both good and poor solvents when the charge density of such PEs exceeds a critical value^17–26^ and with the onset of Manning condensation ^17,27,28^.

There have been a number of efforts to understand the collapse dynamics of neutral polymers by theory, experiment and computer simulation^29–34^. Various simulation studies postulate three or more kinetic stages along the collapse trajectory-an initial stage corresponding to formation and growth of locally collapsed clusters, a coarsening stage characterized by the growth of cluster and a final stage of relaxation. Recently, finite-size analysis has been employed to characterize the coarsening dynamics and aging along the collapsed trajectories of neutral homopolymers^35,36^. A two-point correlation function involving the densities of the local collapsed structures has been proposed to show that the process of aging occurs in collapse of the neutral polymers and a scaling law related to the growth of clusters in the coarsening stage has been proposed as well. While these studies aid in enhancing our insight into the collapse mechanism of the polymers, inclusion of charges along the polymer backbone renders the problem more realistic as the related biopolymers like proteins, DNA and RNA are inherently charged PEs.

Also the inclusion of electrostatic interactions can significantly effect the coarsening dynamics in the PEs and that in turn can aid in cooperative folding from non-native intermediate to native conformation rather than sequential folding by stabilizing energetically favorable tertiary interactions^37^. Long-range electrostatic interactions between sequentially well-separated base pairs in RNA molecules have been suggested to be the basis of formation of motifs such as pseudo knots and associated loop structures whose diverse functional roles are still being investigated^38–40^. Theoretical models and experiments show that the collapsed non-native intermediates play a crucial role in the final folding process rather that the specificity of the initial collapse. Thus folding of biopolymers is a subtle trade-off between conformational entropy of both PE and counterions, electrostatic interactions, solvent effects etc.,^41^. In the previous studies on neutral polymers, quench temperature has been used to reach different end points of “conformational landscape”. In the case of charged PEs, the charge density along the PE plays a similar role in going from extended state (at low charge density) to a collapse state (at high charge densities). While the extended to globule transition of a uncharged polymer chain usually takes place under worsening of the solvent quality^42^, the presence of counterions in the charged polymer systems can additionally influence the collapsed regimes and time scales as has been shown before via different theories^43–45^.

It has been demonstrated in a number of studies that a flexible PE in good solvent undergoes a collapse transition from an extended state with increasing the charge density of the PE. However, in poor solvent, the PE first undergoes a collapsed to extended conformation at low charge densities and further extended to collapse conformation at high charge densities of the PE. Essentially, at very high charge densities of the PE, the solvent conditions play no role in the chain conformation, very unlike the neutral polymer system. In charged polymer systems like PE, the presence of oppositely charged counterions and the balance between the counterion entropy and the favorable electrostatic interactions between the counterions and the charge polymer play a crucial role in the collapse trajectroy. The condensed counterions onto the backbone of the PE, after a critical charge density at a given temperature due to Manning condensation, renormalizes the PE charge and initiates the collapse of the PE^46–50^. In our previous work, we have successfully showed that such condensed counterions can alter the second virial coefficient through their size, further underpinning the role of counterions in the development of effective attractive interactions between similarly charged parts of the PE^49^. All these aspects make studying the kinetics and dynamics of PE, as a function of electrostatic interaction and presence of counterions of different valencies, along the collapse trajectory very relevant. Along with the counterion valency and counterion size, the PE conformation and the mechanism of PE collapse are often highly sensitive to various physiological conditions such as PH, salt concentration, temperature etc. owing to the presence of charge on the polymer chain. The experimental evidence of Ref. 51 and previous numerical simulations.^52,53^ have reported distinctively different responses of a flexible polymer to the presence of various multivalent counterions. In addition, the change in mobility and conductivity of counterions in PE solution with salt concentration has been documented in literature^54^. A salt-induced collapse and re-expansion of highly charged flexible PEs has been observed in Ref. 54 and has been explained by overcompensation of the bare charge of PE. In another recent work of ours, we demonstrated that the aggregation of charged peptides in water and under salt conditions crucially depends on a similar renormalization of the bare charges by the added salt^55^.

To the best of our knowledge the most of the studies on the counterion mediated and salt-induced PE collapse have been focused on the equilibrium properties of PE conformations. In this paper, our aim is to elucidate the kinetic aspects of PE collapse in a poor solvent due to the presence of counterions and further we investigate the effect of monovalent salt concentration on the kinetics of PE collapse. To this end, we have simulated PEs of various length and with varying charge density along the PE backbone in poor solvent conditions. The remainder of the paper of the paper is organized as follows. In Section II we describe the model and methods. In the Section III we present the results of non-equilibrium dynamics related to cluster growth in PE collapse transitions with no additional salt of valency=1,2 and 3. Section IV includes the result of the non-equilibrium studies with the presence of monovalent salt at different concentration and counterions of different valency and Section V contains discussions and conclusions.

## II. MODEL AND METHODS

We model a flexible PE chain as *N* monomers of charge +*qe* (*e* is the elementary charge) connected by harmonic springs,

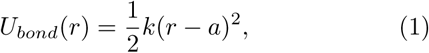

where *k* is the spring constant, *a* is the equilibrium bond length and *r* is the distance between the bonded monomers. The chain and *N*_*c*_=*N/Z* neutralizing counterions, each of charge −*Zqe* where *Z*=1, 2, and 3 for monovalent, divalent, and trivalent counterions, respectively, are placed in a cubic box of linear size *L*. Along with counterions, additional monovalent salt ions are also added in the simulation box. For the case where *N* is not integral multiple of *Z, N*_*c*_ is rounded to closest integer to *N/Z* and then one extra counterion of charge −*qe*(*N* − *ZN*_*c*_) was added to the system to make neutralized system. Thus, for salt-free system, two types of particles exist: monomers and counterions; for systems with additional salt, two additional type of particles are present:monovalent cations and anions of added salt. We express the salt concentration by a term *ρ* where 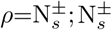 being the number of salt cations/anions of charge ±*e* for monovalent salt *i*.*e* 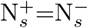. In our simulations the size of all the particles is assumed to be the same and the particles interact via both the excluded volume as well as electrostatic interactions. The volume interaction (van der Waals interactions) between the all pairs non-bonded particles (counterions, salt ions and monomers) separated by a distance *r* is modelled by the 6 − 12 Lennard Jones potential with a cutoff at *r*_*c*_:

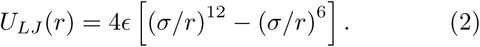

where *ϵ* is the minimum of the potential and *σ* is the interparticle distance at which the potential becomes zero. The electrostatic energy between charges *q*_*i*_ and *q*_*j*_ separated by *r*_*ij*_ is

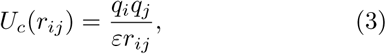

where *ε* is the dielectric permittivity of the solution, taken to be 1 in the present study. The charge density along the PE chain is parameterized by a dimensionless quantity A:

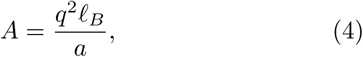

where *ℓ*_*B*_ is Bjerrum length^17^. The Bjerrum length (*ℓ*_*B*_) is the inherent length scale of the charged systems which determines whether electrostatic interaction or the thermal fluctuation dominates and is expressed as,

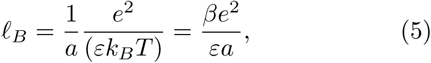

where *k*_*B*_ is the Boltzmann constant, *T* is temperature and *β*=(*k*_*B*_*T*)^−1^. Larger *ℓ*_*B*_ corresponds to higher charge density of the PE.

All the simulations are carried out by molecular dynamics simulation package LAMMPS^56^. The time step for integrating equations of motion is chosen as 0.001 similar to earlier simulations. In the simulations, we use *a*=1.12*σ, k*=500.0 *ϵ*_0_*/σ*^2^, the box size *L* is large enough to guarantee that the PE chain in bad solvent does not act with its own periodic images and the temperature, *k*_*B*_*T/ϵ*_0_=1, is maintained through a Nosé-Hoover thermostat^58,59^. The long-range Coulomb interactions are evaluated using the particle-particle/particle-mesh (PPPM) technique, e.g.^20,60^. We model the poor solvent condition by choosing the following combination of the LJ energy parameters *ϵ*_*LJ*_ and cutoff distance *r*_*c*_: for the monomermonomer interactions:*ϵ*_*LJ*_ =1.5 and *r*_*c*_=2.5. For all other volume interactions among counterions, salt-ions, monomers-counterions and monomers-salt ions, the LJ interactions are repulsive, *ϵ*_*LJ*_ =1.0 and *r*_*c*_=1.0. In current study, multiple systems are considered with a variety of parameters such as charge density (*A*), valency of the counterions (*Z*), PE length (*N*), and salt concentration (*ρ*) of the system and the details of the systems simulated are given in Table I. Most of the analyses are performed for *N*=400 with additional simulations for finite-size scaling analysis. Previous simulation of PE showed a strong evidence of the presence of first-order phase transition from extended to collapse phase, in poor solvent conditions, at *N*=400 as the charge density exceeds a threshold value, consistent with theoretical prediction^20^. Below *N*=400, the first order phase transition is not obvious and this has been attributed to the finite size effects ruling out any other intermediate equilibrium states. For each combination of *A, Z, ρ* and *N*, the initial configuration of the PE is randomly chosen from a short equilibration run such that the counterions are uniformly distributed inside the simulation box and then five independent simulations are performed for each set of parameters and for 2× 10^7^ steps per simulation.

**TABLE I.** The different values of valency of counterions (*Z*), range of charge density of PE (*A*), number of monomers in a PE (*N*) and salt concentration (*ρ*) used in the MD simulations.

| $Z$ | $A$ | $N$ | $\rho$ |
| --- | --- | --- | --- |
| 1 | 0.05-27 | 400 | 0, 100, 200, 400, 600, 800, 1000 |
| 2 | 0.05-12.55 | 400 | 0, 100, 200, 400, 600, 800, 1000 |
| 3 | 0.05-8.03 | 400 | 0, 100, 200, 400, 600, 800, 1000 |

### 1. Cluster identification

To study the non-equilibrium dynamics of the polymer collapse we calculate the average size of monomer clusters *C*_*S*_(*t*), formed in the collapse process. Two non-bonded monomers are considered to be inside the same cluster if the distance between such monomer is less than 2*σ*. The average cluster size is calculated as

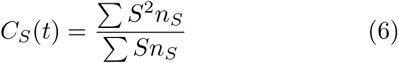

where size of the cluster (*S*) is defined as the number of monomers (*n*_*S*_) within a cluster. We also apply the constraint that the number of monomers in the cluster must be more than 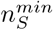 which we have chosen as 10. However, in the long time limit the essential results are not effected for a reasonable choice of 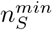.

### 2. Two-time autocorrelation function

To investigate whether the aging phenomenon that was observed in the neutral polymer case can be observed in the charged polymer collapse, we calculate the two-time autocorrelation function *C*(*t*_*w*_, *t* + *t*_*w*_)^61^, defined as,

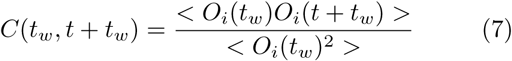

where *O*_*i*_ is a microscopic observable, *t*_*w*_ is the waiting time and the < – – – > denotes the thermal averaging. Here *O*_*i*_ is a microscopic observable, which depends on space 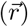 and time (*t*), is chosen in such a way that it characterizes the non-equilibrium changes of any system of interest with time. In case of polymer collapse, we define the observable *O*_*i*_(*t*) at time *t* as a variable depending on the local density around a monomer along the chain which is obtained from the size of the cluster (*S*) to which *i*^*th*^ monomer belongs, defined above. The *O*_*i*_(*r, t*) is assigned as 1 if the monomer is free or outside a cluster and otherwise it is 0. We note that, the starting point (*t*_*w*_) for this analysis is always chosen such that *O*(*t*_*w*_) > 0, i.e., the polymer is in an extended state. Thus estimated *C*(*t, t* +*t*_*w*_) correlates the common density-density correlation function for multi-particle system.

## III. COLLAPSE DYNAMICS OF A PE IN POOR SOLVENT: NO ADDITIONAL SALT

In this section, we investigate the collapse of a PE chain in a poor solvent, neutralized with counterions of various valencies in the absence of additional salt, with the goal of understanding kinetics of such a collapse transition. First, we identify the critical charge density at which the extended to collapse transition occurs (*A*_*c*_) as that determines the range of *A* values to be focussed on. For identifying *A*_*c*_, we monitor the mean square radius of gyration 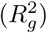 of the PE chain as a function of *A* for counterions of different valencies and the results are shown in Figure 1.

**FIG. 1.**
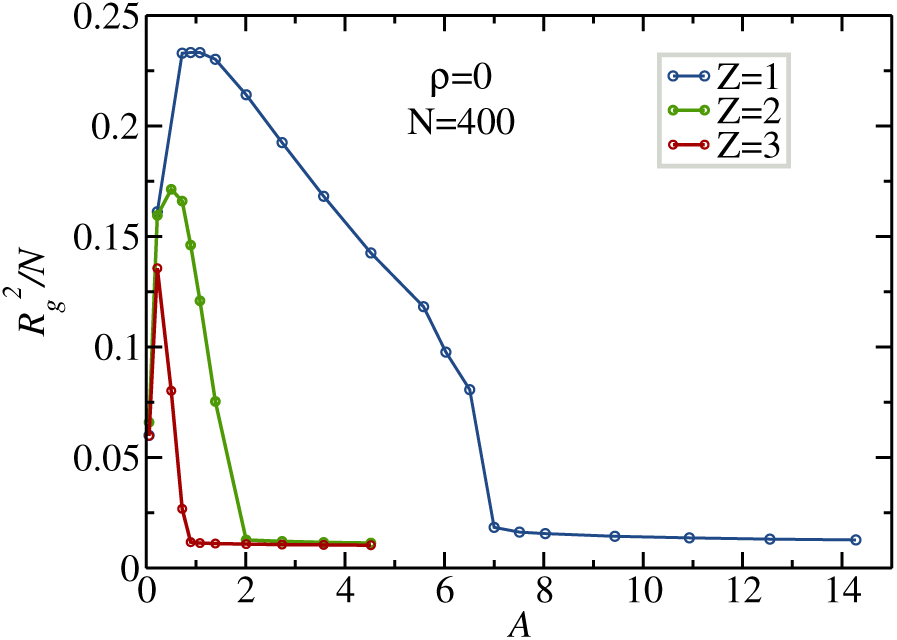
Ratio of the mean square radius of gyration 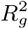 to the no. of monomer *N* as a function of *A* and valency of counterions for a PE of chain length 400 with the presence of only counterions (no additional salt hence *ρ*=0). Each data point is a block average of last 10^6^ steps of production run for five different initial conditions.

Figure 1 shows that under poor solvent conditions, the PE chain adopts a collapsed conformation at very low *A* values (as it is essentially a neutral polymer) and undergoes a collapsed to extended conformation as the value of *A* is increased due to increasing repulsive interactions between the the similarly charged monomers. Until a critical value of *A* is achieved (*A*_*c*_), the counterions occupy the entire volume of the simulation cell, far from the PE chain, to maximize their entropy. For values of *A* greater than *A*_*c*_, the electrostatic interaction between the PE monomers and counterions becomes more attractive compared to the thermal energy of the counterions. This results in the onset of Manning condensation^9,10^ where the oppositely charged counterions begin to condense onto the PE chain causing the similarly charged

PE chain to undergo a counterintuitive extended-collapse transition^17–26^. In our analysis, we have considered the PE chain to be in a collapsed state when the value of 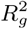 becomes ∼ 0.015*σ* and the critical charge density *A*_*c*_ values are ≈ 8.03, 2.01, 0.893 for monovalent, divalent and trivalent counterions respectively, consistent with our earlier results ^20,60^ for similar chain lengths. The charge of PE monomers corresponding to *A*=8.03, 2.01, 0.893 are 3.00, 1.50 and 1.00 respectively and this suggests that the collapse transition of a PE chain is initiated when the effective charge of PE monomers, *q* ≳3.00/*Z* where *Z* is the valency of the counterion, satisfies *A*_*c*_*Z*^2^ ∼ 9*l*_*B*_*/a*.

### A. Conformational landscape of PE configurations during collapse

In Figure 2 we present the time evolution of the conformational landscape of the PE chain of length *N* = 400 for three different valencies of counterions along the collapse trajectory. The collapsing of PE initiates with the formation of many small clusters connected by linear segments monomers analogous to the pearl-necklace phase described in the literature^43,48,62–67^. The clusters then absorb the remaining monomers, resulting in the compaction of the polymer to form very few large clusters along the PE chain, typically at the two ends of the PE chain connected by bridge of monomers, reminiscent of the dumbbell-like configuration^68,69^. Before final collapse of the PE chain, the few large clusters initiate the coalescence by withdrawing monomers from the bridges leading a sausage-like conformation. The final completely collapsed phase is the result of the internal rearrangement of the sausage-like conformation. Although the sequence of collapse events looks similar, visually, for all *Z* values, the different kinetic stages of formation of nucleation sites of collapse and consequent time scales of collapse transition depend on the valency of the counterion significantly. In order to investigate the kinetics of the PE collapse, we plot the mean square radius of gyration 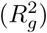 as a function of time for different value of *N* for all the valencies considered in this study (Figure 3). The value of 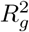 is calculated for values of *A* near the collapse transition (*A*_*c*_).

**FIG. 2.**
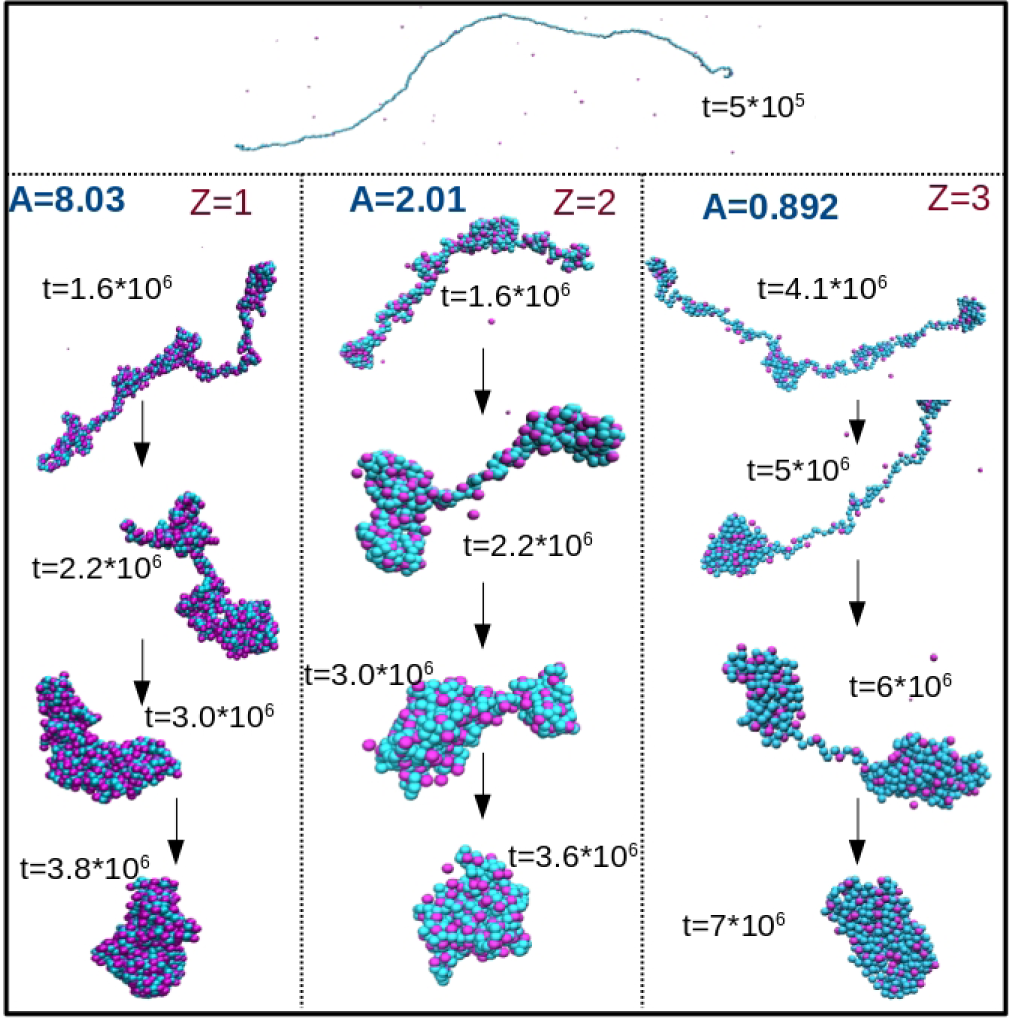
Snapshots of time evolution the polyelectrolyte chain conformations for *Z* = 1, 2, 3 during collapse with *N* = 400. The monomers are shown in cyan and counterions are shown in magenta colors. For the sake of clarity the figure are not shown in same scale.

**FIG. 3.**
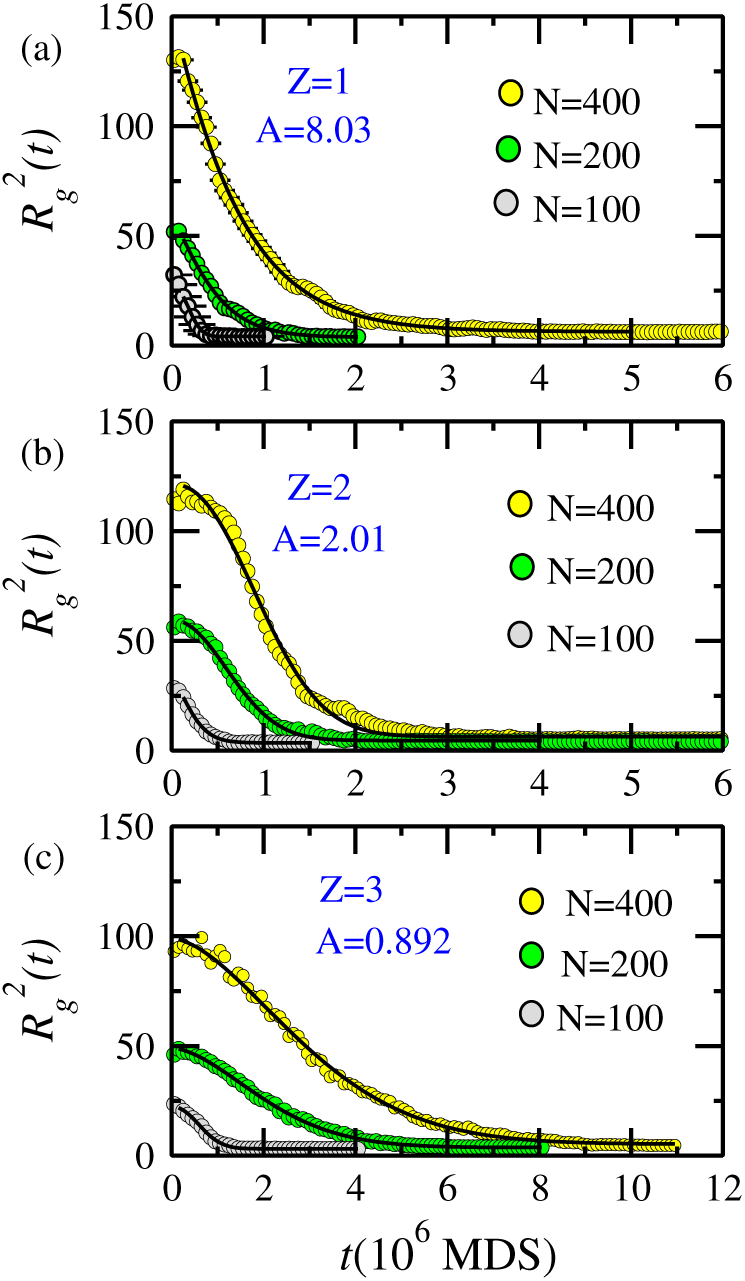
Variation of the mean square radius of gyration 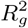 with time near the collapse transition. The circles denotes the original data block-averaged over 100 steps for *Z*=1,2 and 200 steps *Z*=3. The solid black lines are obtained by fitting the original data with the exponential decay: 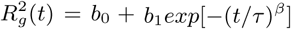.

The variation of 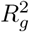 with time step for different values of *N* can be fit to an exponential function of the form:

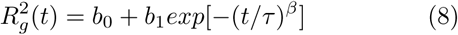

where *b*_0_ corresponds to the value of 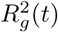 in the collapsed state and *b*_1_ and *β* are fitting parameters. By fitting the curve of 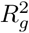 vs *t* we can extract the relaxation time *τ* as shown in Figure 3. Though the exact analytical form of is not known to us, in Figure 3 the solid lines are the corresponding best fits using Equation 8 which captures the decay of 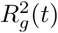 pretty well. Some particular values of the exponents for different chain length *N* at values of *A* very close to collapse transition (*A*_*c*_) are given in Table II and the values of *τ* and *β* for other values of *A* around critical charge density for a PE chain of *N*=400 are listed in Table III for comparison.

**TABLE II.** Numerical values of the exponents obtained from the fit of change of the radius of gyration squared during the collapse of charged polymer of different chain length *N*. Each value was obtained by averaging data from 5 trajectories of the system.

| $Z$ | $A$ | $N$ | $\tau$ ($10^3 \times$ MDS) | $\beta$ | $N$ | $\tau$ ($10^3 \times$ MDS) | $\beta$ |
| --- | --- | --- | --- | --- | --- | --- | --- |
| 1 | 8.03 | 100 | 203.31 | 1.77 | 370 | 783.34 | 1.22 |
|  |  | 200 | 503.6 | 1.37 | 400 | 764.14 | 1.09 |
|  |  | 250 | 425.31 | 1.60 | 500 | 858.12 | 0.12 |
|  |  | 270 | 675.44 | 1.34 | 600 | 963.18 | 0.19 |
|  |  | 300 | 560.77 | 1.28 | 700 | 1045.81 | 0.44 |
|  |  | 350 | 656.57 | 1.6 | 800 | 1148.12 | 0.75 |
| 2 | 2.01 | 100 | 288.12 | 1.74 | 370 | 1192.0 | 2.28 |
|  |  | 200 | 855.74 | 1.94 | 400 | 1217.48 | 2.08 |
|  |  | 250 | 619.33 | 2.02 | 500 | 927.44 | 1.26 |
|  |  | 270 | 1052.32 | 1.68 | 600 | 1238.1 | 1.59 |
|  |  | 300 | 876.07 | 1.63 | 700 | 2088.2 | 0.31 |
|  |  | 350 | 1066.61 | 2.36 | 800 | 3577.4 | 0.43 |
| 3 | 0.893 | 100 | 767.80 | 2.11 | 370 | 3525.95 | 1.74 |
|  |  | 200 | 2460.56 | 1.77 | 400 | 4019.1 | 1.88 |
|  |  | 250 | 1787.12 | 1.78 | 500 | 4478.8 | 1.95 |
|  |  | 270 | 3118.46 | 1.75 | 600 | 4987.1 | 1.90 |
|  |  | 300 | 2426.31 | 1.93 | 700 | 5873.7 | 2.35 |
|  |  | 350 | 2959.92 | 2.00 | 800 | 6907.5 | 1.99 |

**TABLE III.** Values of parameters *τ* and *β* of the kinetic law in Equation 8 for PE chain of *N*=400 at different *A* related to collapse transition.

| $Z$ | $A$ | $\tau$ ($10^3 \times \text{MDS}$) | $\beta$ |
| --- | --- | --- | --- |
| 1 | 7.51 | 817.23 | 0.076 |
|  | 8.03 | 764.14 | 1.09 |
|  | 9.43 | 766.55 | 1.35 |
|  | 14.28 | 649.39 | 1.95 |
| 2 | 1.75 | 1657.51 | 1.27 |
|  | 2.01 | 1217.48 | 2.08 |
|  | 2.73 | 645.16 | 1.63 |
|  | 4.51 | 456.35 | 1.90 |
| 3 | 0.72 | 5990.41 | 1.77 |
|  | 0.893 | 3525.95 | 1.74 |
|  | 1.39 | 1226.52 | 3.04 |
|  | 4.51 | 348.47 | 2.13 |

From the estimated *τ* at different *N* we can predict the exponents (*ν*) relating the relaxation time to the chain length (*τ* ∼ *N*^*ν*^) for the physical process responsible for the collapse kinetics of PE. Kuznetsov et al^33^ in their kinetic studies on the collapse transition of neutral homopolymer by Monte Carlo simulation performed without considering the hydrodynamic effects explained the kinetics by three stage processes: formation of clusters, growth of local clusters and the final coarsening stage where the final coarsening stage was expressed by the law 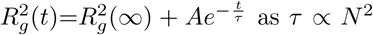. Further the same group studied the collapse neutral homopolymers quenched from a good to poor solvent in dilute solution using Gaussian self-consistent approach without hydrodynamics and confirmed their previous argument about the scaling laws of the characteristic time of the coarsening stage *i*.*e. τ* ∼ *N* ^2^)^70^. Also, in the same work they showed that the value of the exponent *ν* changed to 3/2 with the inclusion of hydrodynamic interaction^70^. The analytical work in Ref. 34 presented a scaling argument describing the kinetics for the polymer collapse where the collapse time varies as *τ*_*H*_ ∼ *N* ^4/3^ and *τ*_*B*_ ∼ *N* ^2^ in hydrodynamics and brownian solvent respectively, and it agreed well their numerical results. In another study^71^ on the polymer collapse using Monte Carlo simulations of a bead-spring model where the relaxation timescale measured in a similar way by Equation 3 was shown to obey rouse scaling behavior *τ* ∼ *N* ^2^;^72^ which comparable with the exponent obtained from Brownian dynamics simulations of bread-spring model of Ref. 34. However, the values predicted in the literature for the exponents *ν* vary from 1.3 to 2. To find the dependence of characteristic timescale of PE collapse on the size of the monomer, *τ* as a function of *N* is plotted in Figure 4. In the study of collpase transition of strongly charged polymers, we find that the finite-size scaling behavior of *τ* doesn’t follow the rouse scaling as observed before in the case neutral polymer^34^,71, instead the dependence of *τ* with *N* in non-monotonic in nature for short length of PE chain. For *N* ≥ 400, *τ* varies as ∼*N* ^0.59^ for *Z* = 1 and for *Z* = 3, *τ* is proportional to *N* ^1.14^ for *N* ≥ 600. But for *Z* = 2, the quality of exponential fitting of 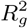 was poor, so we have not obtained any monotonic dependence of *τ* on *N* but the the value of *ν* seems to lie as 0.59 < *ν* < 1.14. We note that the values of exponent *ν* in PE collapse is less than those of neutral polymer collapse regardless of valency of counterions. This suggests the significance of long-range electrostatic interactions in the collapse dynamics and *τ* increases with the increase of size of PE chain regardless of the valency of counterions. From Table III, it can be seen that for *A* values *A* > *A*_*c*_, there is a consistent decrease in the relaxation times for all valencies of the counterions as the PE chain enters a strongly collapsed regime. It can also be seen from Table II that for the same chain length of *N* = 400, the relaxation time increases with the valency of the counterions strongly suggesting that the transition of PE chain from an extended state to a collapsed conformation is slower in the presence of multivalent counterions. Now we define the collapse time (*t*_*c*_) as the time when 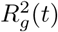 has decayed to 90% of its total decay, *i*.*e*., 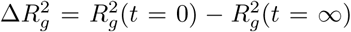. More quantitatively we can say at *t* = *t*_*c*_, 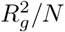 falls approximately to 0.035*σ*-0.032*σ* for *N*=400. The *t*_*c*_ of a PE molecule is the time at which PE chain starts to collapse from its intermediate compact states, can be estimated by fitting parameter of Equation 8. The *t*_*c*_ of PE chain for different length is listed in Table IV. The Table IV shows that *t*_*c*_ increases with the increase of *N* regardless the value of *Z*. So we find that in the case of charged polymer the location of the collapse point is very much dependent on charge parameter *A* and the valency of counterions.

**FIG. 4.**
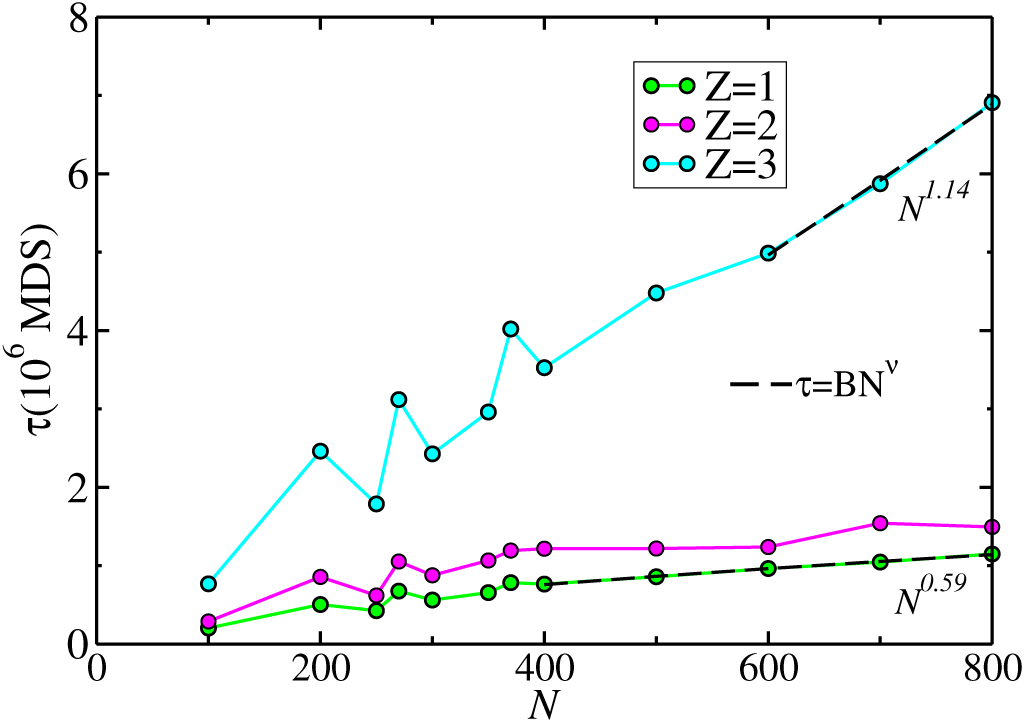
Scaling of the relaxation times *τ* at around *A*_*c*_ for different valency of counterions, estimated from Equation 8, as a function of the chain length. The dashed lines show show the respective fit as *τ* = *BN* ^*ν*^ obtained by using the data for *N ⩾* 400.

**TABLE IV.** The variation of collapse time with degree of polymerization *N*.

| $Z$ | $A$ | $t_c^{N=100}$ | $t_c^{N=200}$ | $t_c^{N=300}$ | $t_c^{N=400}$ |
| --- | --- | --- | --- | --- | --- |
| 1 | 8.03 | $3.38 \times 10^5$ | $9.41 \times 10^5$ | $1.08 \times 10^6$ | $1.82 \times 10^6$ |
| 2 | 2.01 | $4.83 \times 10^5$ | $1.23 \times 10^6$ | $1.48 \times 10^6$ | $2.79 \times 10^6$ |
| 3 | 0.893 | $1.18 \times 10^6$ | $3.36 \times 10^6$ | $3.79 \times 10^6$ | $5.58 \times 10^6$ |

We now calculate the time evolution of the cluster size states (*S*_*i*_) of each monomer of the PE chain in poor solvent condition for different valencies of the counterions and for values of *A* close to *A*_*c*_, is measured via the location of a monomer within a cluster of size *n*_*S*_ as shown in Figure 5. We identified the cluster size state (*S*_*i*_(*t*)) of each monomer at each time step and we have shown it in a color plot as a function of time in Figure 5. The cluster size state (*S*_*i*_(*t*)) of the *i*^*th*^ monomer at time *t* is the size of the cluster to which it belongs at time *t*. The plot shows that at intermediate times, large clusters coexist with small clusters for all counterion valencies. However, for monovalent counterions the collapse of entire PE chain into one large cluster occurs at much earlier times compared to the counterions of other valencies. The plot also clearly demonstrates that for trivalent counterions, the formation of a single cluster occurs at much later times, consistent with earlier experimental results^73,74^.

**FIG. 5.**
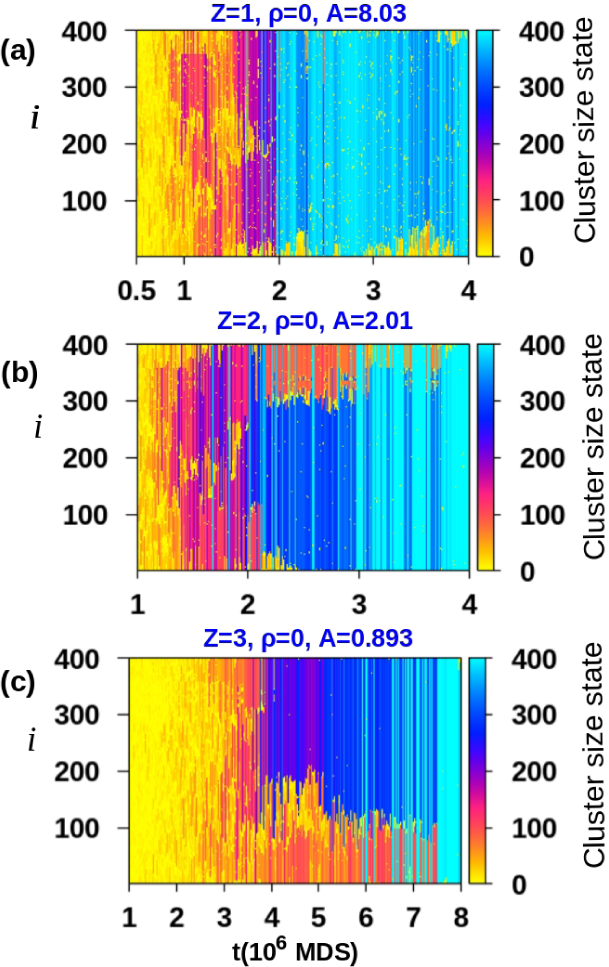
Time evolution of cluster size state (*S*_*i*_(*t*)) of monomers along the length of polymer with *N*=400 for different valency of counterions where corresponding *A* values are around *A*_*c*_ associated to different valency of counterions.

To further quantify the dependence of the time evolution of average cluster size of a PE chain, along the collapse trajectory, on the valency of counterions, the mean cluster size (*C*_*S*_(*t*)) as a function of time is measured and the results are shown in Figure 6(a-c). Figure 6(a-c) shows a comparison of cluster growth timescale between monovalent, divalent and trivalent counterions for a fixed length (*N* =100, 200, 400) of PE chain. As can be seen from the data of Figure 6, the coarsening of the cluster formation is significantly different for trivalent counterions and the chain length also plays an important role. In the case of trivalent counterions we also observe a delay in the nucleation of small clusters during the non-equilibrium process compared to other two cases which is shown in Figure 6(d) as a zoomed version of Figure 6(c) at the initial stage of cluster formation.

**FIG. 6.**
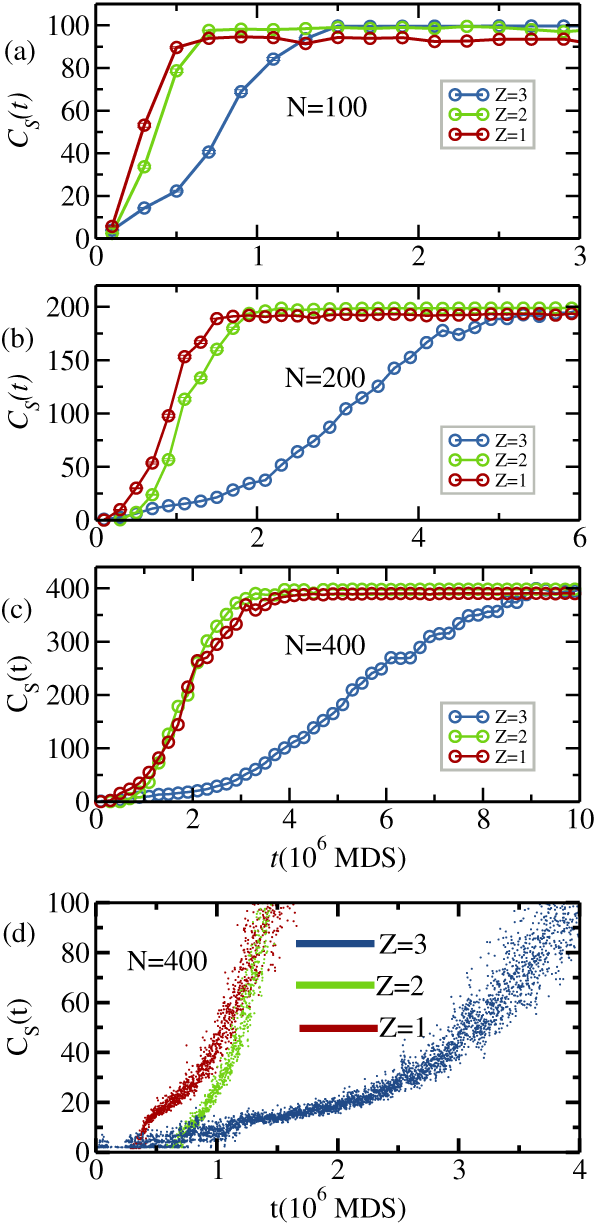
(a-c)Comparison of the growth of the average cluster size *C*_*S*_(t) with time for a polymer of fixed length *N* =100, 200, 400 at the *A*=8.03, 2.01, 0.893 for *Z*=1,2, 3 respectively. Here *A* values correspond to the critical charge density with no salt for three different valency of counterions. (d)Zooming in of *C*_*S*_(*t*) vs *t* plot at initial stage of clusters formation for *N*=400.

The collapse timescale of the PE chain for monovalent and divalent counterions is very similar for values of *A* larger than *A*_*c*_, though for values of *A* ∼ *A*_*c*_, there are considerable differences in the collapse time scales. This can be attributed to the difficulty in determining the exact value of *A*_*c*_ for PE systems. Keeping this in mind, in Figure 7 we have shown *C*_*S*_ *vs t* at three different *A* that leads to collapse transitions and compared with that of *A* (*A* < *A*_*c*_) for which final conformation is not the collapsed one. In Figure 7 the solid lines are the fitting of the data corresponding to scaling behaviour of coarsening which is discussed in later section (Sec. III C). We have observed that slight tuning of *A* at larger values fastens the collapse time and the collapse transition is almost instantaneous at sufficiently large *A i*.*e*. there is a jump from collapse state to globule state and is independent on the valency of counterions.

**FIG. 7.**
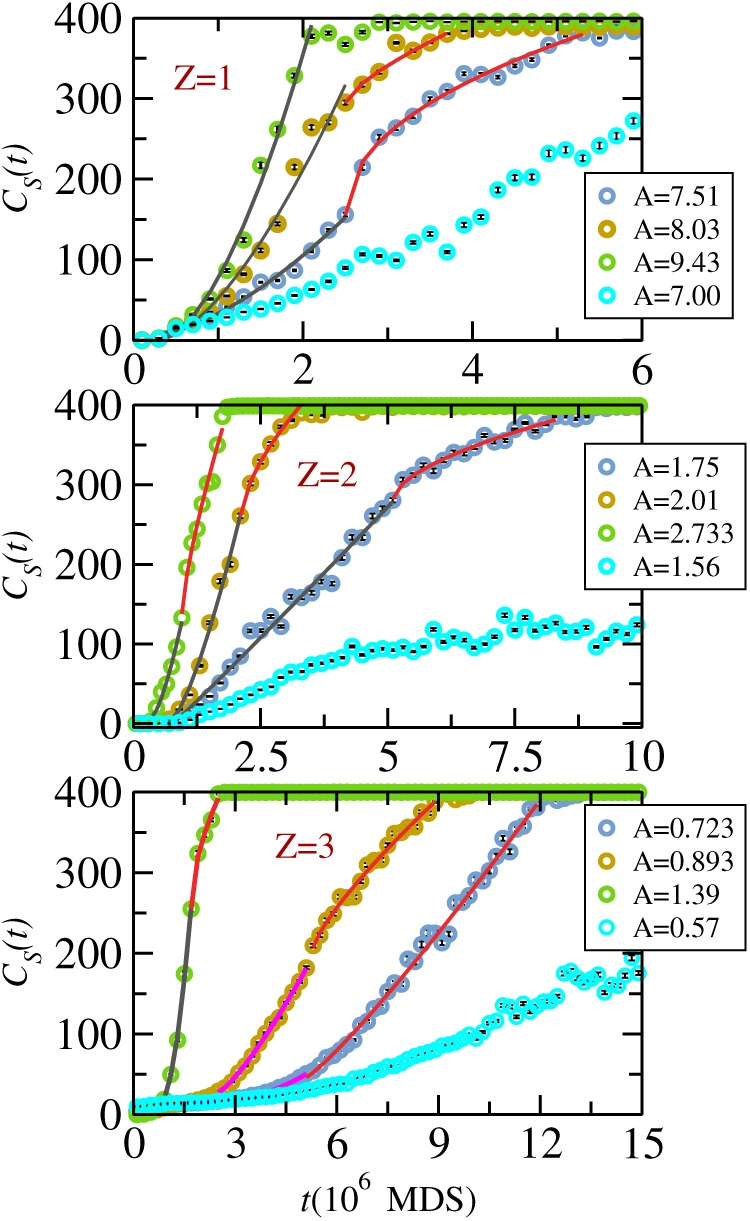
Average cluster size (*C*_*S*_(t)) as a function of *t* at different *A*.

### B. Time evolution of condensed counterions

To explore the role played by the condensation of the counterions in the observed difference in the onset of collapse as a function of valency, we calculate the number condensed counterions (*n*_*cond*_) as a function of time. In our analysis, a counterion is considered to be condensed if its distance from any monomer is less than 2*σ*. The mean fraction of condensed counterions as a function of time for all the counterion valencies considered in the study is shown in Figure 8. For the values of *A* considered here, which is around the critical value required for the Manning condensation to occur, the rise in the number of condensed counterions is very rapid for the all counterions. However the data strongly suggests that for the same length of the PE chain, the onset of complete counterion condensation onto the PE chain is significantly different for different valencies of the counterions. Specifically for divalent and trivalent counterions, even at longer simulation times compared to *Z*=1 valency, all the counterions do not condense on the chain and only for *A* >> *A*_*c*_ all counterions were found to be condensed (data is not shown here) on PE chain, strongly suggesting that the complete condensation of counterions is not necessary for collapse transition. It should be noted that for *N* = 400 and trivalent counterions, one extra counterion of charge −*qe* was added along with 133 counterions of charge −3*qe*. We measure *t*_*cond*_ as the time when atleast 90% of the counterions are condensed and the values of *t*_*cond*_ for monovalent, divalent and trivalent counterions with *N*=400 are 9.2 × 10^5^ at *A*=8.03, 5.5 × 10^6^ at *A*=2.01 and 8.4 × 10^6^ at *A*=0.893 respectively. The difference in the counterion condensation times seen here is strongly correlated with the difference in the collpase times of the PE chain as well as the time evolution of the cluster sizes. It is to be noted that though the effective charges on the counterions are different depending on the valencies, the van der Waals radius of the counterions remains the same. This will result in lesser number of multivalent counterions inside the collapsing phase of the PE molecule as compared to the monovalent counterions and effective shielding of the PE monomer charges may be more effective in the case of monovalent counterions in terms of effective distances between similarly charged monomers. This difference may be more pronounced at the charge density of the PE increases and any small fluctuations in the condensed counterion positions can affect the effective attractive interactions in the otherwise similarly charged system significantly.

**FIG. 8.**
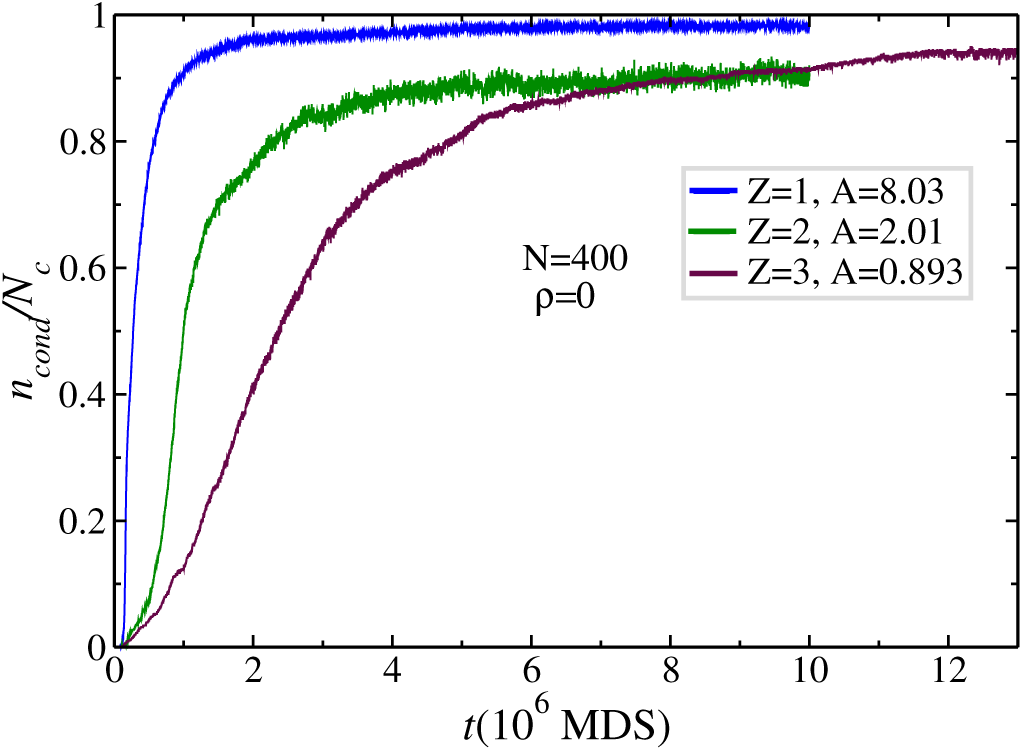
Mean fraction of counterions * *n*_*cond*_ ∗*/N*_*c*_ > within a distance 2*l*_*B*_ from the PE chain (*N* = 400) as a function of time for *Z*=1,2 and 3 and the corresponding *A* values are 8.03, 2.01 and 0.893 respectively.

### C. The scaling behaviour of coarsening

In the case of neutral polymers, it was observed that there are two regimes of growth of clusters: the initial cluster formation is slower and scales as *C*_*S*_ ∼ *t*^1/3^ and beyond a crossover, the cluster growth is faster and scales as *C*_*S*_ ∼ *t*^0.98^ (Ref. 71). To explore the presence of similar collapse rates in the case of charged polymers like PE, the scaling behavior of cluster growth along the collapse trajectory near *A*_*c*_ values for all the valencies is shown on a double-log scale in Figure 9. The data clearly shows that, in contrast to the neutral polymer case described above, the cluster growth rate in the case of PE system is faster at earlier times and slower at later times and there is evidence of multiple time scales along the collapse trajectory as a function of valency of counterions. This is reminiscent of coarsening stage and the *C*_*S*_(*t*) grows with time *t* as a power law, *C*_*S*_(*t*) ∼ *t*^*α*^ with different exponents for earlier and later stage of PE collapse. The time-scale of crossover from faster growth to slower growth stage is also different for different valencies (*Z*) of the counterions. In Figure 9, we fitted the raw data as *C*_*S*_(*t*) = *C*_*S*_ (*t*_0_) + *B*(*t* − *t*_0_)^*α*^ instead of *C*_*S*_(*t*) = *Bt*^*α*^ to incorporate different initial points for subsequent stages of cluster growth. We find that the dynamical exponent, *α*, is a function of the valency of counterions and *A*. The chosen value of *A* in Figure 9 is close to *A*_*c*_ as it would be difficult to find exact critical value of *A* for collapse transition. From our simulation results we might expect the critical value of *A* as 7.51 ≤ *A*_*c*_ ≤ 8.03, 1.75 ≤ *A*_*c*_≤ 2.01 and 0.723 ≤ *A*_*c*_ ≤ 0.893, for *Z*=1,2 and 3 respectively. The varying values of dynamic exponent with different *Z* is in complete contrast to the unique value of of *α* in collapse transition of neutral polymer^71^ strongly suggesting the role of electrostatic interactions and in particular the effective renormalization of charges of PE by the counterions. The fitting parameter of cluster growth size at different *A* corresponding to collapse transition is listed in Table V and the corresponding fitted data at *A* ≈ *A*_*c*_ are shown in Figure 9 by solid lines. We find that while the collapse trajectory is a two-stage kinetic process for monovalent and divalent counterions, it is a three-stage kinetic process for trivalent counterions and can be seen from the three different cluster growth regimes in Figure 9 for trivalent counterions. It is also to be noted that very high charge density values of PE chain (*A* >> *A*_*c*_), the coil-globule transition is almost instantaneous (see Figure 7) and this suggests the need to carefully select the range of *A* values over which such coarsening behavior can be observed.

**TABLE V.** Results from the fitting of the average cluster size *C*_*S*_(*t*) vs *t* with the form *C*_*S*_(*t*) = *C*_*S*_ (*t*_0_) + *B*(*t* − *t*_0_)^*α*^ with three different interaction parameter *A* corresponds to collapse transition.

| $Z$ | $A$ | kinetic stage | $t_0(10^6 \text{ MDS})$ | $t_f(10^6 \text{ MDS})$ | $C_S(t_0)$ | $B$ | $\alpha$ |
| --- | --- | --- | --- | --- | --- | --- | --- |
| 1 | 7.51 | initial | 0.3 | 2.5 | 1.75 | 0.001 | 1.55 |
|  |  | final | 2.5 | 5.3 | 155.12 | 5.74 | 0.46 |
|  | 8.03 | initial | 0.3 | 2.5 | 2.09 | 0.001 | 1.57 |
|  |  | final | 2.5 | 3.7 | 294 | 0.56 | 0.71 |
|  | 9.43 | single | 0.3 | 2.1 | 2.37 | 0.001 | 1.72 |
| 2 | 1.75 | initial | 0.7 | 5.1 | 1.08 | 0.05 | 1.03 |
|  |  | final | 5.1 | 8.3 | 280.5 | 0.8 | 0.59 |
|  | 2.01 | initial | 0.7 | 2.1 | 5.72 | 0.01 | 1.41 |
|  |  | final | 2.1 | 3.1 | 260.83 | 1.4 | 0.65 |
|  | 2.733 | initial | 0.3 | 1 | 3.9 | 0.01 | 1.44 |
|  |  | final | 1 | 1.8 | 132 | 1.53 | 0.75 |
| 3 | 0.723 | initial | 0.9 | 2.7 | 1.59 | $7.1 \times 10^{-6}$ | 0.99 |
| | | intermediate | 2.7 | 5.1 | 18.09 | $5.03 \times 10^{-3}$ | 1.96 |
| | | final | 5.1 | 11.9 | 50.78 | $1.42 \times 10^{-2}$ | 1.14 |
|  | 0.893 | initial | 0.9 | 2.3 | 8.33 | 0.005 | 1.09 |
|  |  | intermediate | 2.3 | 5.1 | 29.01 | 0.0076 | 1.23 |
|  |  | final | 5.1 | 8.9 | 181.97 | 0.59 | 0.71 |
|  | 1.39 | initial | 0.7 | 1.7 | 8.66 | 0.0002 | 1.99 |
|  |  | final | 1.7 | 2.5 | 255 | 3.5 | 0.54 |

**FIG. 9.**
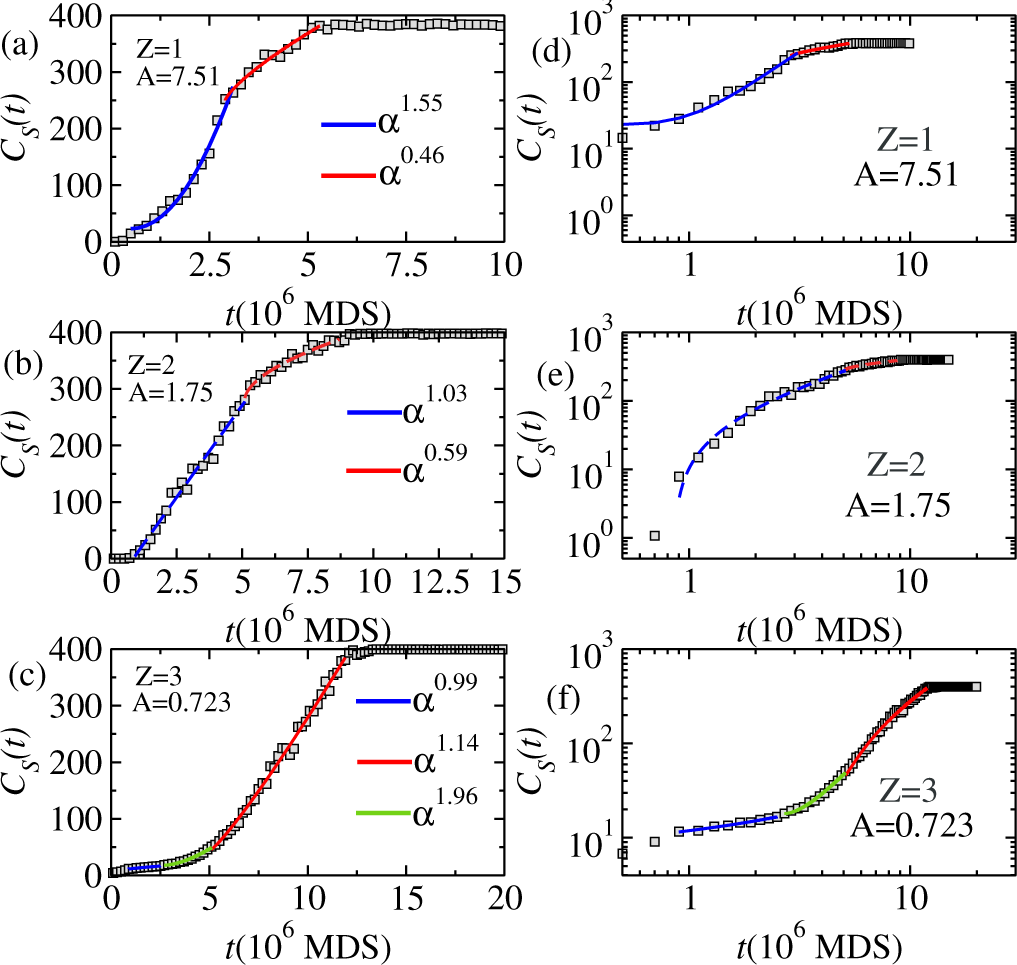
(a-c) The average cluster size *C*_*S*_(*t*) versus *t* during the coarsening kinetic stage for different valency of counterions. The solid lines are a fit to the form *C*_*S*_(*t*) = *C*_*S*_ (*t*_0_) + *B*(*t* − *t*_0_)^*α*^ where *t*_0_ is the initial point of a kinetic stage. (d-f) The same data on a double-log scale.

### D. Aging dynamics in collapse of PE

Earlier simulations on neutral polymer collapse^29,30,33,34,70,71,75–77^, using Monte Carlo, Gaussian self-consistent methods or Langevin approaches have shed light on our understanding of the kinetics of the polymer collapse transition. Another important aspect for our deeper understanding of such non-equilibrium processes is the aging property^61,78–81^. This phenomenon, which is well known experimentally in structural (e.g., polymer) glasses captures the non-equilibrium slow dynamics processes, can be examined by calculating the two-time autocorrelation function (*C*(*t*_*w*_, *t* + *t*_*w*_)) involving the waiting time *t*_*w*_ and the observation time *t*. For aging dynamics, the larger the waiting time, the longer the system takes to forget the initial configuration. The presence of aging is displayed in two-point autocorrelation function if there is an absence of translation invariance in such function as well as a slowing down of relaxation with increasing the waiting time *t*_*w*_ occurs. Two time correlation functions, (*C*(*t*_*w*_, *t* + *t*_*w*_)) characterising the non-equilibrium kinetics, were measured for all the three valencies and results are shown in Figure 10(a-c). In Figure. 10 the autocorrelation function for different waiting times, as a function of *t* − *t*_*w*_ is shown. Included in the inset is the data as a function of *t/t*_*w*_ on double-log plot for Z=1,2 and 3 respectively for PE chain of length *N* = 400. In the *C*(*t*_*w*_, *t* + *t*_*w*_) vs *t* − *t*_*w*_ plots, no violation of time-translational invariance was observed. Further one can easily extract relaxation time (*t*_*r*_) associated with the decay of *C*(*t*_*w*_, *t* + *t*_*w*_) and this relaxation time will be a rapidly increasing function of *t*_*w*_ if there is a presence of aging. By visual inspection of the inset plots of Figure 9 one could look the decrease in relaxation time with increase of *t*_*w*_. Therefore, no presence of aging is observed in the collapse transition of charged polymer in contrast a flexible single neutral polymer collapse by quenching from extended to globular phase as reported in Ref. 71. Aging is observed in non-equilibrium systems with slow dynamics, whereas the condensation counterions stimulates the coil-globule transition and so, the non-equilibrium dynamics of PE collapse does not share the phenomenology of aging.

**FIG. 10.**
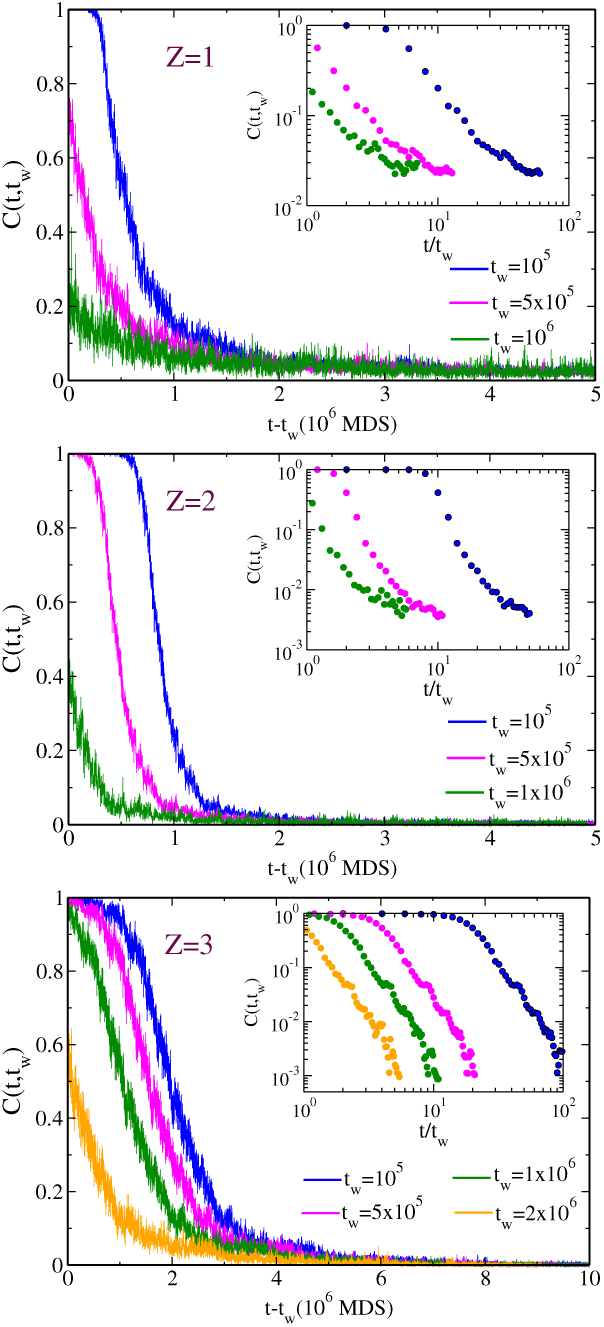
Autocorrelation function *C*(*t, t*_*w*_) as a function of *t* − *t*_*w*_ for different waiting times for PE with *N*=400. Inset shows simple scaling with respect to *t/t*_*w*_ on a double-log scale. The corresponding *A* values are *A*=8.03, 2.01 and 0.893 for *Z*=1, 2 and 3 respectively.

### E. Long-range contact formation of during PE collapse

The initial globule or sausage conformation (as defined in Section III A) has many intramolecular knots or contacts which are relaxed via reptation of the crumpled globule. A macromolecule must undergo compaction from a disordered extended conformation to a native structure to be in its functional form. Compact (or collapsed) intermediates, which can be formed above critical charge density, provide a context for long-distance tertiary interactions which eventually help to speed up the collapse process. Formation of such tertiary contacts can influence the cooperativity of base pairing and the process of helix assembly via collapse in case of RNA/DNA folding^8,82^ and hence the tertiary contacts form a crucial bridge between the amorphous collapsed phase and the final native state.

A recent study shows multivalent cations stabilize compact forms of the RNA more efficiently than monovalent ions^83^. To find the impact of counterion valency on the long range interaction we calculate the number of long-range contacts (*N*_*nl*_) as a function of time (Figure 11) for different valencies of counterions. In this calculation any contact between the *i*^*th*^ and *j*^*th*^ monomer is considered to be a long range contact if the monomer index difference *i*−*j* ≳50 and are at a distance of 2*σ* from each other. The data shown in Figure 11 shows that though the onset of long-range tertiary contacts in the case of trivalent counterions is later than for counterions of lower valencies, the number of contacts is much higher. The data also suggests that the initiation of tertiary contacts is earliest in the case of monovalent counterions, the the number of contacts is an increasing function of valency of the counterions. A visual representation of collapsed state of PE chain formed at *t*=1.4 × 10^7^ time steps in the presence of counterions of *Z*=1, 2,and 3 are shown in Figure 12. In Figure 12 we can see the compaction of PE chain is more in the presence of trivalent counterions than other two cases that in turn may enhances the chances of formation of tertiary contacts that might aid the folding of system towards the native conformation.

**FIG. 11.**
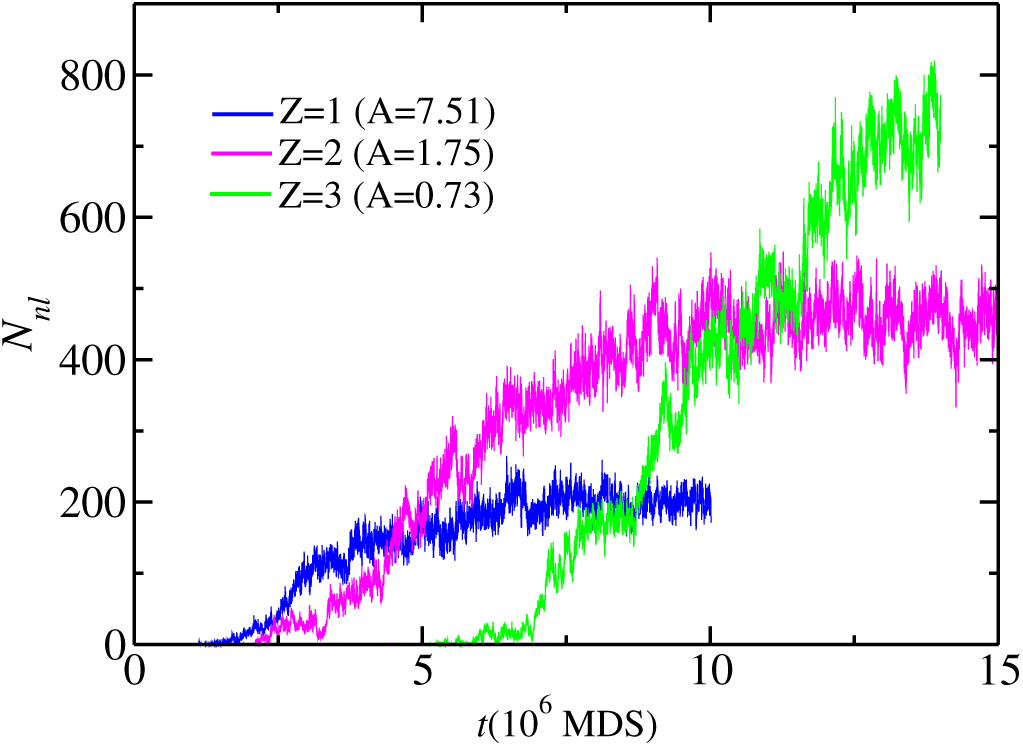
Number non-local long-range contacts (*N*_*nl*_) as a function of time of a PE chain of size *N* = 400 for different valencies of counterions.

**FIG. 12.**
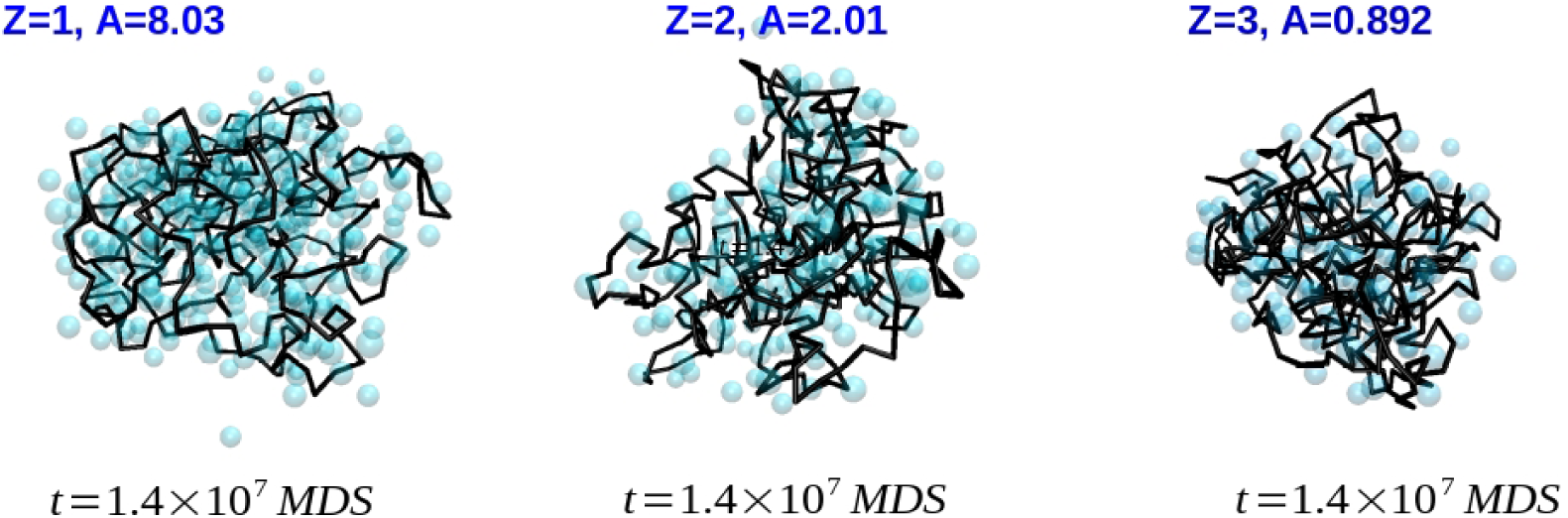
Snapshots of the polyelectrolyte chain conformation at time steps 1.4 × 10^7^ for different valency of counterions. The black solid line shows the PE conformation and the counterions are shown by light-cyan color circles.

## IV. NON-EQUILIBRIUM DYNAMICS OF PE COLLAPSE:EFFECT OF SALT CONCENTRATION

### A. Identification of collapse point at salt concentration

Next we investigate how the presence of additional monovalent salt ions along with counterions affects the kinetics of PE collapse and the intermediate configurations of PE chain. We first determine the value of *A*_*c*_ at various salt concentration for three different valencies of counterions and for this, we monitor the 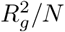 of the PE chain as a function of charge density in the presence of salt at various concentrations and compare with the control case of no additional salt (*ρ* = 0) (Figure 13). The data strongly suggests the following: (a) the presence of additional salt delays the occurrence of complete collapse phase towards higher values of *A* for all counterion valencies (b) there is also evidence that the presence of salt can change the nature of transition from extended to collapsed phase. Our earlier simulations^20^ and other works have suggested that the transition from extended to collapse transition is first-order in nature without an intermediate thermodynamic phases. The data in Figure 13 shows that the enhanced entropic contribution of salt ions and excluded volume interaction between counterions and salt-ions may cause a decrease in effective interactions of monomer-counterions and monomer-salt ions delaying and/or changing the nature of the phase transition. The increase in *A*_*c*_ with the increase of *ρ* might be due to the opposition to counterions-monomers interaction by enhanced counterions-salt-ions interaction.

**FIG. 13.**
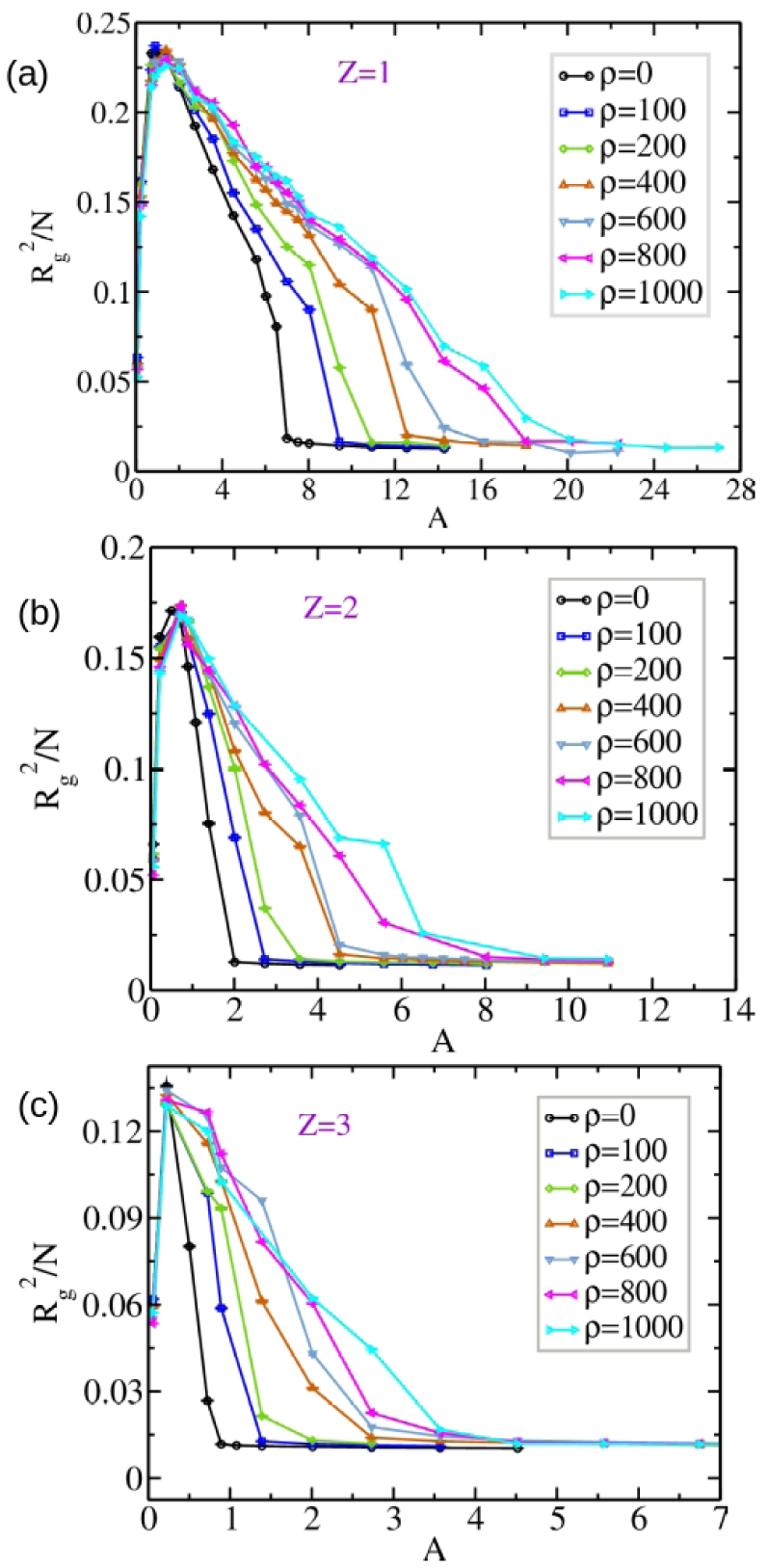
Ratio of the mean square radius of gyration 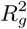 to the number of monomers *N* = 400 as a function of *A* and for different salt concentrations. Each data point is an average of 5 number of simulations.

To understand the effect of inclusion of additional salt ions on the critical charge density of collapse (*A*_*c*_), we list the values of *A*_*c*_ for all chain lengths (*N*) considered, for three different counterion valencies and for three different salt concentrations in Table VI. The table suggests the following trends: (1) For chain lengths and valencies, there is an increase in the values of *A*_*c*_ with the addition of additional salt ions (2) for shorter chains, the *A*_*c*_ values seem to saturate for higher salt concentrations. It is to be noted that accurate determination of *A*_*c*_ is difficult and the values listed in the Table VI are best within the limitations of the MD simulations at values of *A* considered here. However, the trends that the *A*_*c*_ values show are rather clear as explained above. The Table VI also suggests that the addition of monovalent salt affects the case of monovalent counterions the most, due to similarity in the size and charge between counterions and salt ions and it would be interesting to probe, in future studies, the effect of different sizes of counterions and salt and their role in competitive collapse transition of the PEs. In case of multivalent counterions, the counterions can exchange with the bound salt monovalent ions, due to higher efficiency in inducing compaction with fewer counterions and this is reflected in the lesser degree to which the values of *A*_*c*_ change with addition of salt in their case.

**TABLE VI.** The critical charge density of PE chain at different salt concentration with PE chain of various length and various valency of counterions.

| Z | $N_m$ | $A_c$<br>($\rho=0$) | $A_c$<br>($\rho=200$) | $A_c$<br>($\rho=400$) | $A_c$<br>($\rho=600$) |
| --- | --- | --- | --- | --- | --- |
| 1 | 400 | 8.03 | 12.55 | 16.12 | 20.14 |
|  | 200 |  | 14.28 | 20.14 | 24.6 |
|  | 100 |  | 16.12 | 20.14 | 24.6 |
|  | 50 |  | 12.55 | 14.28 | 14.28 |
| 2 | 400 | 2.01 | 3.57 | 4.52 | 5.58 |
|  | 200 |  | 4.52 | 6.75 | 6.75 |
|  | 100 |  | 4.52 | 5.58 | 5.58 |
|  | 50 |  | 2.733 | 3.57 | 3.57 |
| 3 | 400 | 0.893 | 2.01 | 2.733 | 3.57 |
|  | 200 |  | 2.733 | 3.57 | 3.57 |
|  | 100 |  | 2.01 | 2.733 | 2.733 |
|  | 50 |  | 0.893 | 0.893 | 0.893 |

### B. Kinetics of PE collapse with various salt concentration

Next we focus our attention to the influence of salt ion on the kinetics of PE collapse. In Figure 14 we plot the the decay of the radius of gyration 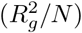 of PE with time for *N*=400 and *Z*=1 at different values of *ρ*. As *ρ* increases, we observe that there are indications that the phase change of PE chain during collapse might change form first order transition towards continuous transition. Also, beyond *ρ* > 600 there is no significant difference in the rate of decay of 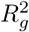. For *ρ* > 600, 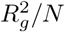 decays to 0.015*σ* at *t* ≈ 10^7^ steps. It suggests that regardless of the size of PE chain, the effect of salt ions tends to saturate when the number of salt ions are sufficiently large.

**FIG. 14.**
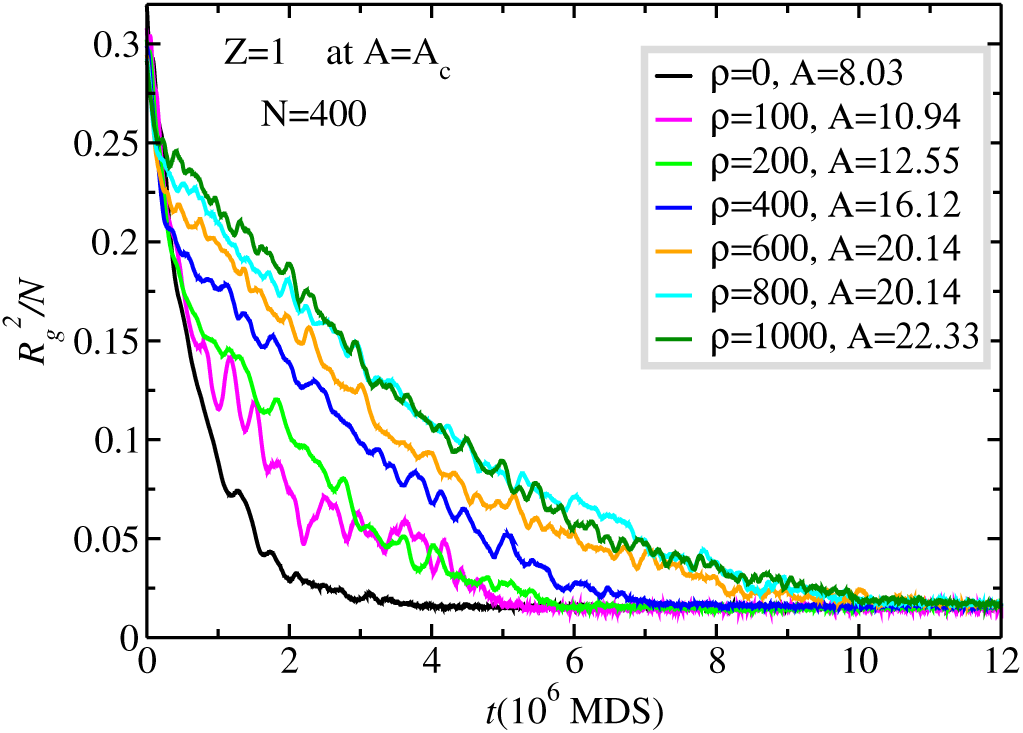
Variation of the ratio of the mean square radius of gyration 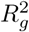 to *N* at *A* = *A*_*c*_ corresponding to varying salt concentration with time during collapse.

Figure 15 shows the growth of cluster size as a function of time at different salt concentration for *N*=400 and the same for four different values of *N* in Figure 16. The data shows that in the case of monovalent and divalent counterions, addition of salt slows down the clustering along the PE collapse trajectory, while in the case of trivalent counterions, the clustering is little faster with addition of salt. It can also be seen that the additional monovalent salt ions effect the clustering dynamics of the PE collapse significantly more when compared to the trivalent counterions, supporting the earlier observations in critical charge density (Table VI). In Figure 16, the coarsening dynamics of the PE collapse as a function of chain length of the PE is shown. In Figure 16(d), (h) and (l) we can see the salt effect on growth rate of *C*_*S*_(*t*) is more pronounced at higher concentration (*ρ* = 600) with *Z*=1, 2 compared to *Z*=3 irrespective the length of PE chain. To understand the cause of relative speed up of the PE collapse in trivalent counterion case with monovalent salt ions we have to look at the PE configuration formed during collapse which has been discussed in next section.

**FIG. 15.**
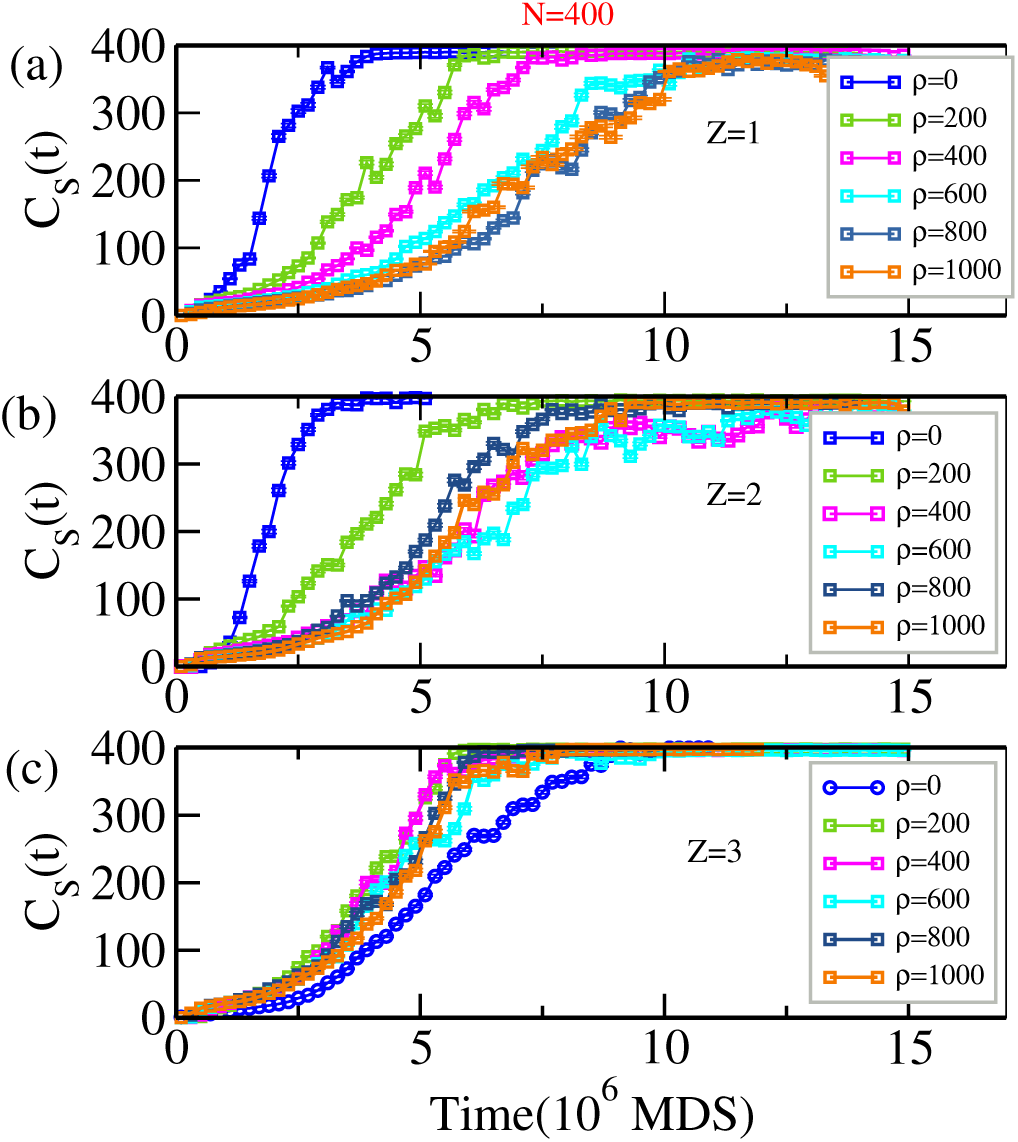
Growth of the average cluster size *C*_*S*_(*t*) with time for a polymer with *N*=400 with different salt concentration *ρ*=0-1000 for *Z*=1,2, 3 respectively. Here *A* values correspond to the critical charge density with additional salt for three different valency of counterions as listed in Table VI.

**FIG. 16.**
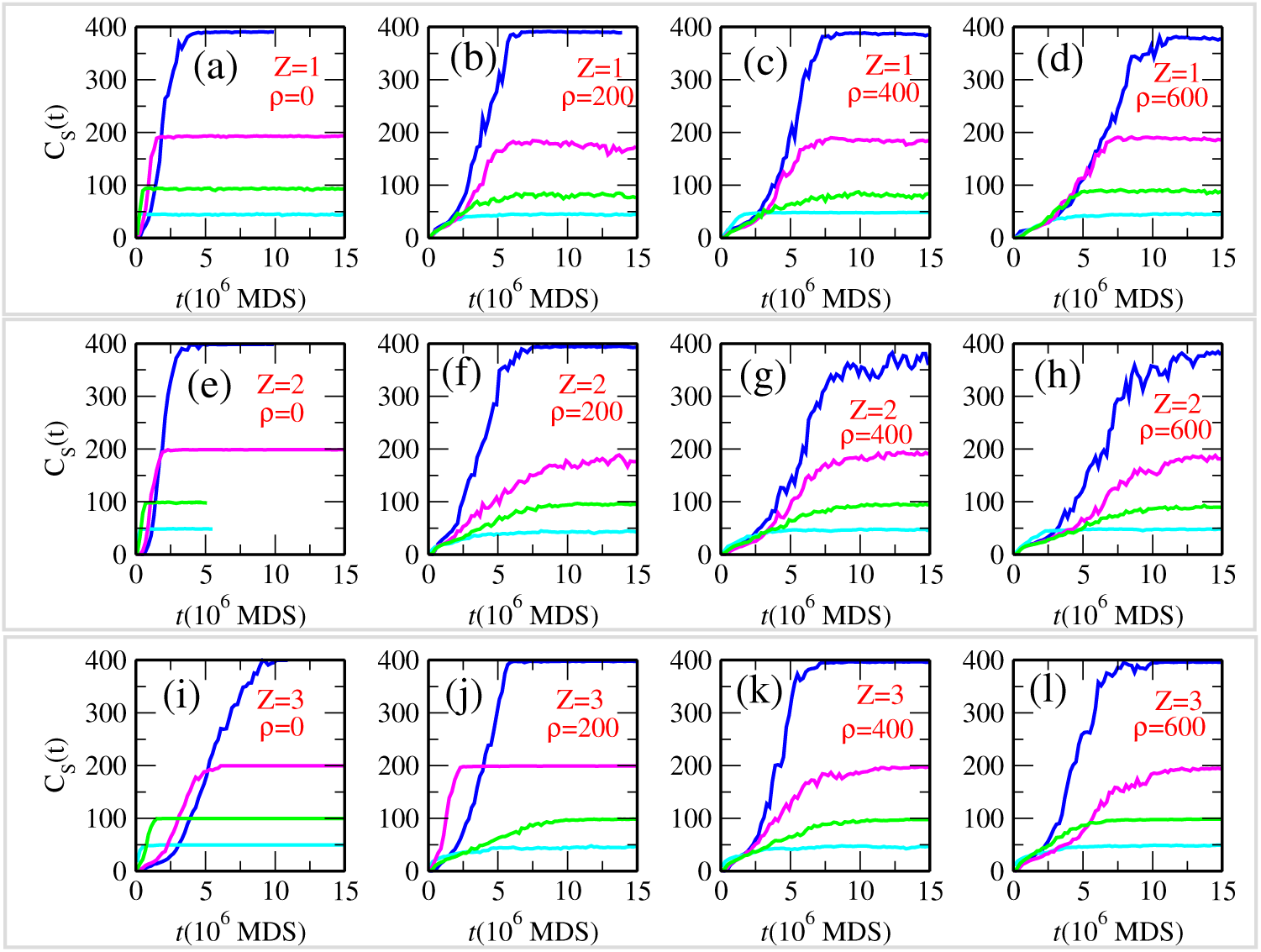
Growth of the average cluster size *C*_*S*_(*t*) with time for a polymer for *N* =50, 100, 200 and 400 with varying salt concentration and varying valency of counterions at their respective *A*_*c*_ values as listed in Table VI. The cyan, green, magenta and blue color denote data for the PE chain of *N* = 50, 100, 200 and 400 monomer respectively.

### C. Effect of salt ions on PE configurations during collapse

For trivalent counterions, we have observed that there are some transient long-range contacts formed frequently with the addition of salt, which could speed up the cluster growth for trivalent counterions. Such snapshots of long range contact formations for *Z*=3 is shown in Figure 17. In the trajectory of PE chain with additional salt and trivalent counterions there were formation of loop like conformation. Also the dumbell like structures was not found profoundly on the trajectory like the case of *ρ*=0; pearl-necklace and sausage like configurations are only the intermediate on the collapsing pathway. The kinetic mechanism of PE collapse with additional salt and trivalent counterions differs from that of without salt case.

**FIG. 17.**
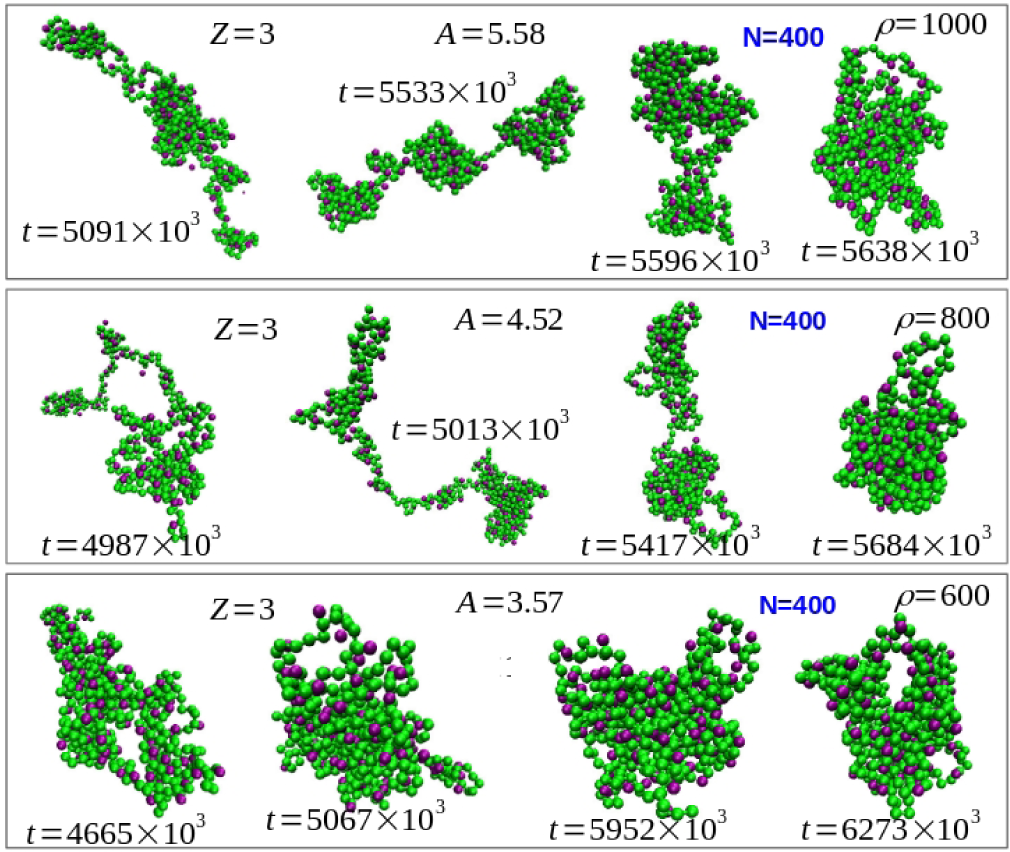
Snapshots of PE configurations observed in one of the trajectories for collapse transition with *ρ*=1000, 800 and 600 and trivalent counterions. Green and purple colors depict the PE-ions and counterions respectively.

Here the predominant effects, at early time involves the nucleation of small clusters, coarsening of small clusters in pearl-necklace structure with small loop formation and results in final collapsed species by a cluster-cluster coarsening mechanism. To quantify the effect of salt on the entanglement of the collapse structure we calculate the number of non-local contacts (*N*_*nl*_) formed as a function of time with additional salt conditions and is compared with control system as shown in Figure 18. In Figure 18 we observe that the presence of additional salt along with counterions suppresses the total number of non-local contact in final collapse structure, regardless the valency of counterions *i*.*e*. the collapsed PE conformation is less entangled in the presence of additional salt. Also, noteworthy that the monovalent salt-ions effect mostly the degree of entanglement of PE chain when *Z*=3 as for *Z*=3, as *N*_*nl*_ dropped from 900 to 500 approximately for *Z*=3. It has been suggested that RNA folding occurs from collapsed non-specifically intermediate states by searching for its tertiary contacts within a highly restricted subset of conformational space^6^. In addition, the over-compaction of a conformational state can slow the folding dynamics by fostering formation of nonnative interactions and by providing steric barriers to formation of the native configuration. Here rapid and comparatively high cooperative collapse of a PE molecule and small loop formation upon addition of salt with trivalent counterions opens up the possibility of rapid folding to native-state.

**FIG. 18.**
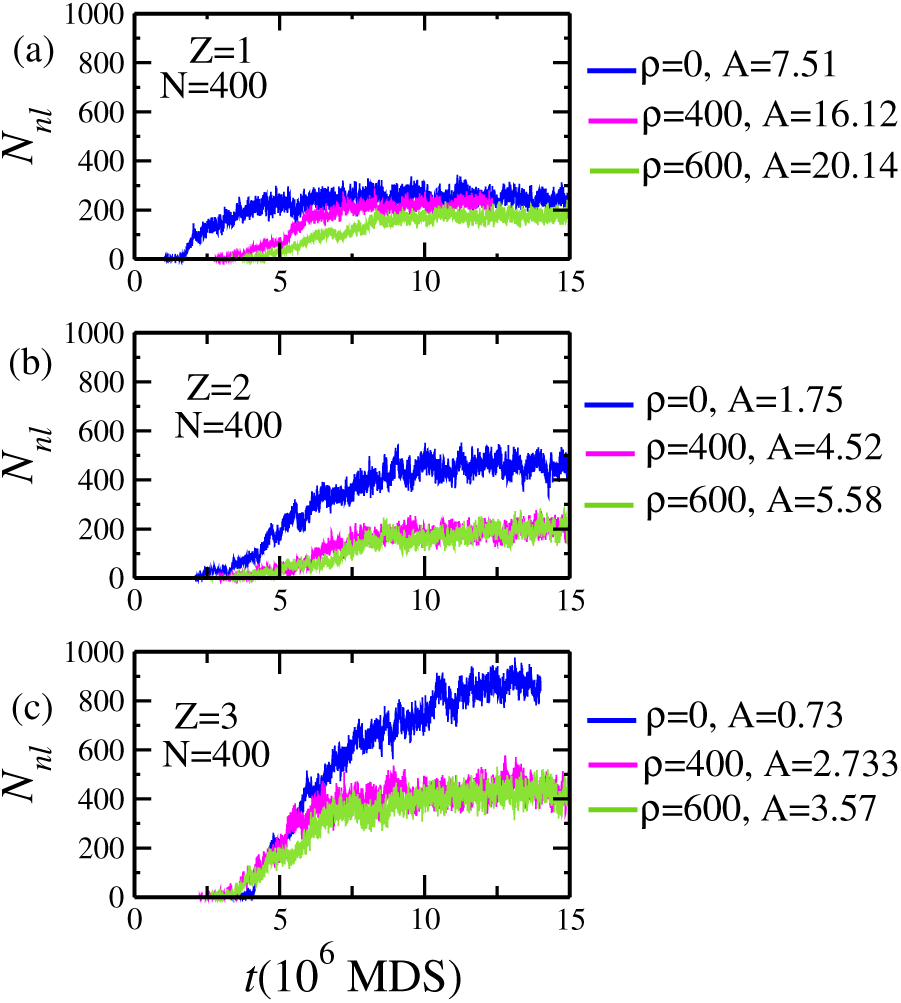
Comparison of number non-local contacts as a function of time of a PE chain of size *N* = 400 between control system (*ρ*=0) and system with additional salt concentration (*ρ*=400, and 600) for (a)*Z*=1 (b)*Z*=2 and (c)*Z*=3.

## V. DISCUSSIONS AND CONCLUSIONS

In this work, we have used molecular dynamics simulations to study the kinetics of flexible polyelectrolyte collapse mediated by counterion condensation and also to find the influence of monovalent salt ions on the kinetics of PE collapse. In our simulations the counterions and salt ions were considered explicitly while the solvent was implicit. We considered the case of a poor solvent for the chain with repulsive interactions for both the counterions and salt-ions. We mainly focused on the dependence of the swelling and the collapse of PE chain on a dimensionless parameter *A*, characterizing the relative strength of electrostatic interaction, the valency *Z* of counterions, the chain length *N* and the salt concentration *ρ*.

In accordance with the previous studies^20,60^, at very low values of the *A* the PE chain adopts a collapse conformation. As *A* is increased the PE chain is swollen up due to the Coulomb repulsion between the chain ions and with the further increase of *A* the 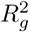 of the chain decreases and finally, at a particular value of *A*, the PE chain undergoes collapse transition due to condensation-induced intra-chain attraction resulting in a shrinkage of the chain. This critical value of *A i*.*e. A*_*c*_ decreases with the increase of counterion valency. Our results generalize a quantity *A*_*c*_*Z*^2^ to hold a constant value. The sequence of events observed during the PE collapse is very similar for all valencies of counterions considered and also very similar to those observed during collapse transition of a neutral polymer^71^. The collapse of polyelectrolyte chain starts by the nucleation of primary clusters separated by linear segments or bridges and the nucleation sites grow in size by accumulating monomers from the bridges which results in the “pearl-neclace” structure^65^, followed by coarsening of small clusters into a dumbell-like structure and eventually the coalescence of clusters to form a collapsed conformation. Though the sequence of events in similar regardless of the valency of counterions, the rate of collapse transition was found to decrease with the increase of valency of counterions. The collapse transitions with monovalent counterions occur 3-fold faster than transitions induced by trivalent counterions for the same size of the PE chain. The condensation of counterions is expected to renormalize the total charge of PE chain and lead to polyelectrolytes to a non-specially collapsed state. The slower rate of condensation of multivalent counterions is responsible for the delay in PE collapse in mutivalent counterions case and we have observed that complete condensations of counterions or complete charge neutralization is not necessary for obtaining a collapsed state. For *A*>*A*_*c*_ the rate of phase transition of PE was found to increase with *A* and for sufficiently large *A* the coil-glouble transition will be almost instantaneous *i*.*e* within a time of 10^6^ steps. The present paper represents an effort to find a finite-size scaling by describing the kinetics by a power law decrease in time of the squared radius of gyration with an exponent. The predicted characteristic times of the coarsening stage in the presence of monovalent (*τ* ∼ *N* ^0.59^) and trivalent (*τ* ∼ *N* ^1.14^) counterions were found to be much less than that of neutral polymer collapse (*τ* ∼ *N* ^4/3^) in hydrodynamic heat bath^84^. Our analysis relating scaling behaviors related to the growth or coarsening of clusters of monomers (*C*_*S*_(*t*) ∼ *t*^*α*^) indicates that the the overall collapse kinetics of a PE system has several underlying distinct kinetic mechanisms that result in two/three kinetic regimes. In our earlier work, we demonstrated different regimes of collapse^49^ and the present results also support that observation. The value of the exponent is found to be the function of parameter *A* and valency of counterions which is unlike the neutral polymer collapse. In addition the evidence of aging dynamics has not been obtained in PE collapse like in the dynamics of polymer collapse^71^. The overall compaction of extended polyelectrolyte chain to a globule shape is highest for trivalent collapse. Such tightly packed structure with multivalent counterions might aid the RNA/DNA folding from non-native collapsed state by providing a platform for the formation of tertiary contacts.

Futher studies on the non-equilibrium dynamics of the polyelectrolyte-counterion system in the presnce additional monovalent salt suggest that critical value of *A* for collapse transition is less sensitive to salt concentration for trivalent counterions and this result is consistent with the result of Ref. 85. The critical value of *A* is increased with the increase of salt concentration for all valency of counterions. In the presence of salt-ions and with monovalent and divalent counterions, the rate of phase transition becomes slower with the increase of salt concentrations and it becomes saturated for sufficiently high salt concentration. Remarkably, the presence of salt-ions slightly speed up the collapse transition for trivalent counterions and they have profound influence on the intermediate states on the collapsing pathway. Observed formation of long-distance loops during the collapse trajectory, especially in the presence of trivalent counterions, is suggested to be the cause of acceleration in phase transition of PE chain with trivalent counterions. Also, a significant decrease in the degree of compactness of final collapsed state is observed for trivalent in the presence of additional monovalent salts.

## Acknowledgments

The simulations were carried out on the supercomputing machines Annapurna and Nandadevi at the Institute of Mathematical Sciences.

